# Long-read sequencing based genomic data of a dipluran species, *Occasjapyx japonicus*

**DOI:** 10.64898/2026.07.16.716246

**Authors:** Kakeru Yokoi, Atsushi Toyoda, Kosei Hashimoto, Tsunaki Asano

## Abstract

We present the genome dataset of a dipluran species, *Occasjapyx japonicus*, representing the first dipluran genome assembled using HiFi long-read sequencing technology. The assembled genome is approximately 439.3 Mbp in size, comparable to those of other dipluran species available in public databases. The N50 value of 15.5 Mbp exceeds that reported for other dipluran species. The assembled gene set contains 19,635 genes, a number not significantly different from those estimated in previous analyses of two other dipluran species. Functional gene annotation was conducted using predicted amino acid sequences derived from the gene set. BUSCO analysis indicated that the assembled genome contains the majority of conserved core genes. These findings suggest that the *O. japonicus* genome and associated data are of sufficient quality to serve as a reference genome. The dataset will be valuable for studies in comparative or evolutionary biology, particularly in understanding hexapod evolution and the emergence of insects.

## Background

Insects belong to the phylum Arthropoda, which is characterized by a segmented body and external skeleton. Recent molecular studies indicate that insects originated from a crustacean lineage (remipedes), challenging the earlier view that they are more closely related to myriapods. Historically, the taxon Insecta included primitive non-winged hexapods such as Collembora, Protura, and Diplura. In contemporary phylogenetics, these three taxa are classified as “non-insect hexapods” distinct from the “true” insects[1,2]. Non-insect hexapods possess three pairs of legs and a tripartite body plan (head, thorax, and abdomen), features also observed in true insects. However, non-insect hexapods are classified as Entognatha, with mandibles enclosed within the head capsules, whereas true insects (Ectagonatha) have exposed mandibles. Among the three groups of non-insect hexapods, diplurans have been regarded as the closest relatives of insects, based on both morphological and molecular evidence[2,3]. Recent analyses have proposed that a taxon, comprising both diplurans and collembolans, forms a sister group to Insecta[4]. Even though, the phylogenetic placement of diplurans (one of the closest arthropod relatives of insects) has garnered interest from entomologists seeking to understand the origins of insects.

The order Diplura is divided into several taxa, including Japygidae and Campodeidae, both of which contribute to its taxonomic diversity[5]. Members of Japygidae are distinguished by predatory tail appendages adapted for grasping. The first publicly available genomic dataset for Diplura was an annotation set generated through the i5K project for *Catajapyx aquilonaris*[6], followed by the genome report of *Campodea augens*[7]. Both genomes were assembled using short-read sequencing, with gene models predicted using RNA-seq data as hints. Additionally, genomic datasets for two more dipluran species, *Campodea plusiochaeta* and *Campodea silvestrii*, are now available in the NCBI genome database (Accession IDs: GCA_034698565.1, and GCA_034694985.1, respectively)[8,9]. For both species, the genome assemblies were generated using short-read genomic data.

Recent advances in DNA sequencing technologies have enabled the generation of long-read-based genome assemblies for numerous non-model species. These high-contiguity genome assemblies facilitate accurate gene prediction, particularly for tandemly arranged genes and improve sequence resolution in repetitive sequences. To our knowledge, no dipluran genome has been assembled using long-read sequencing technologies. Here, we present the genome dataset of the dipluran species *Occasjapyx japonicus. O. japonicus* belongs to the suborder Dicellurata, one of the two dipluran suborders (Rhabdura and Dicellurata), characterized by the presence of tail forceps (Fig. 1)[5].. To the best of our knowledge, our data set in this study is the first dipluran genome assembled using HiFi long-read sequencing.

**Fig. 1.**
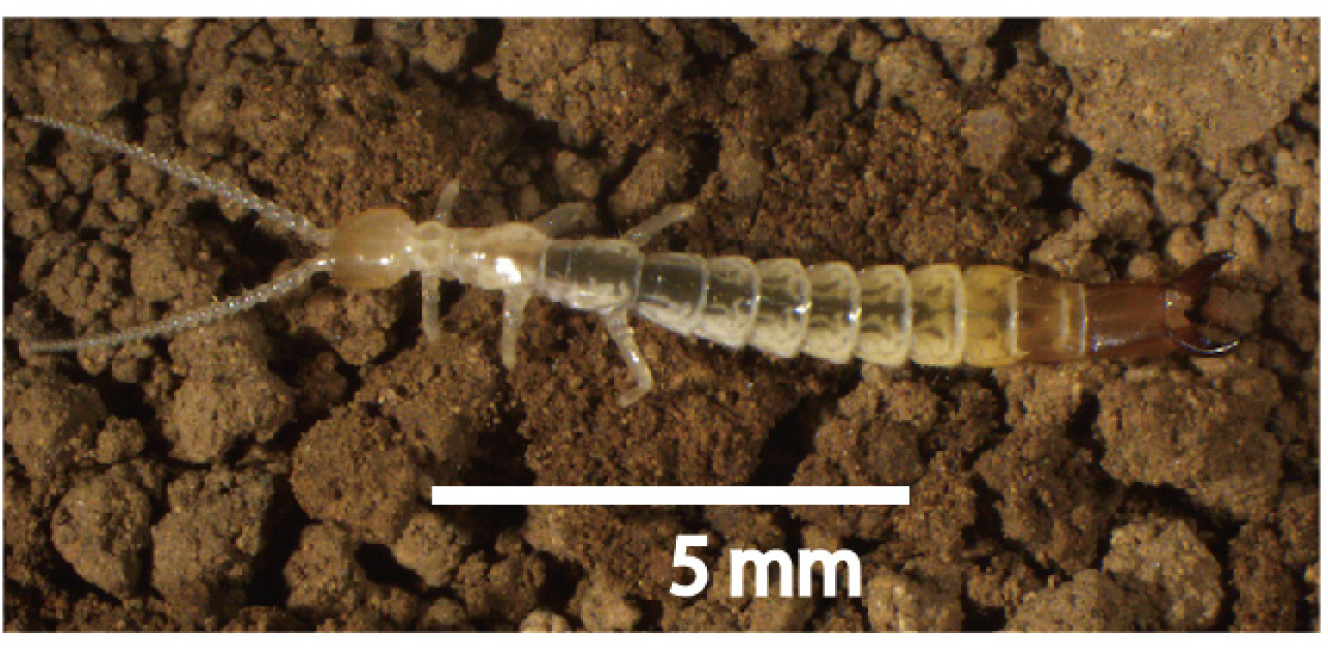
Photograph of the forceptail, *Occasjapyx. japonicus*.

## Methods

### Sequencing procedures

Specimens of the forceptail, *O. japonicus* (Fig. 1), were collected from the campus of Tokyo Metropolitan University (35°37⍰16.7⍰N 139°23⍰5.5⍰E), and identified based on the descriptions of Enderlein (1907), Kuwayama (1928) or Silvestri (1928)[10–12] supplemented by more recent information from Pagés (1989)[13].

Total RNA was extracted individually from each whole-body specimen using the NucleoSpin RNA plus XS kit (Takara Bio, Tokyo, Japan). The extracted RNA was used for cDNA library preparation with the SMRTbell Prep kit 3.0 (Pacific Biosciences, CA, USA). PCR amplification and purification of the cDNA were performed according to the manufacturer’s protocols. The concentration and the size distribution of each library were measured using the Qubit 4 Fluorometer (Thermo Fisher Scientific, MA, USA) and Agilent 2100 Bioanalyzer (Agilent Technologies, CA, USA), respectively. Each library was sequenced on the PacBio Sequel II platform using one Sequel II SMRT Cell 8M with the Sequel II Binding Kit 3.2 and Sequencing Kit 2.0 (Pacific Biosciences, CA, USA); the movie collection time was 30 hours (Fig. 2).

**Fig. 2.**
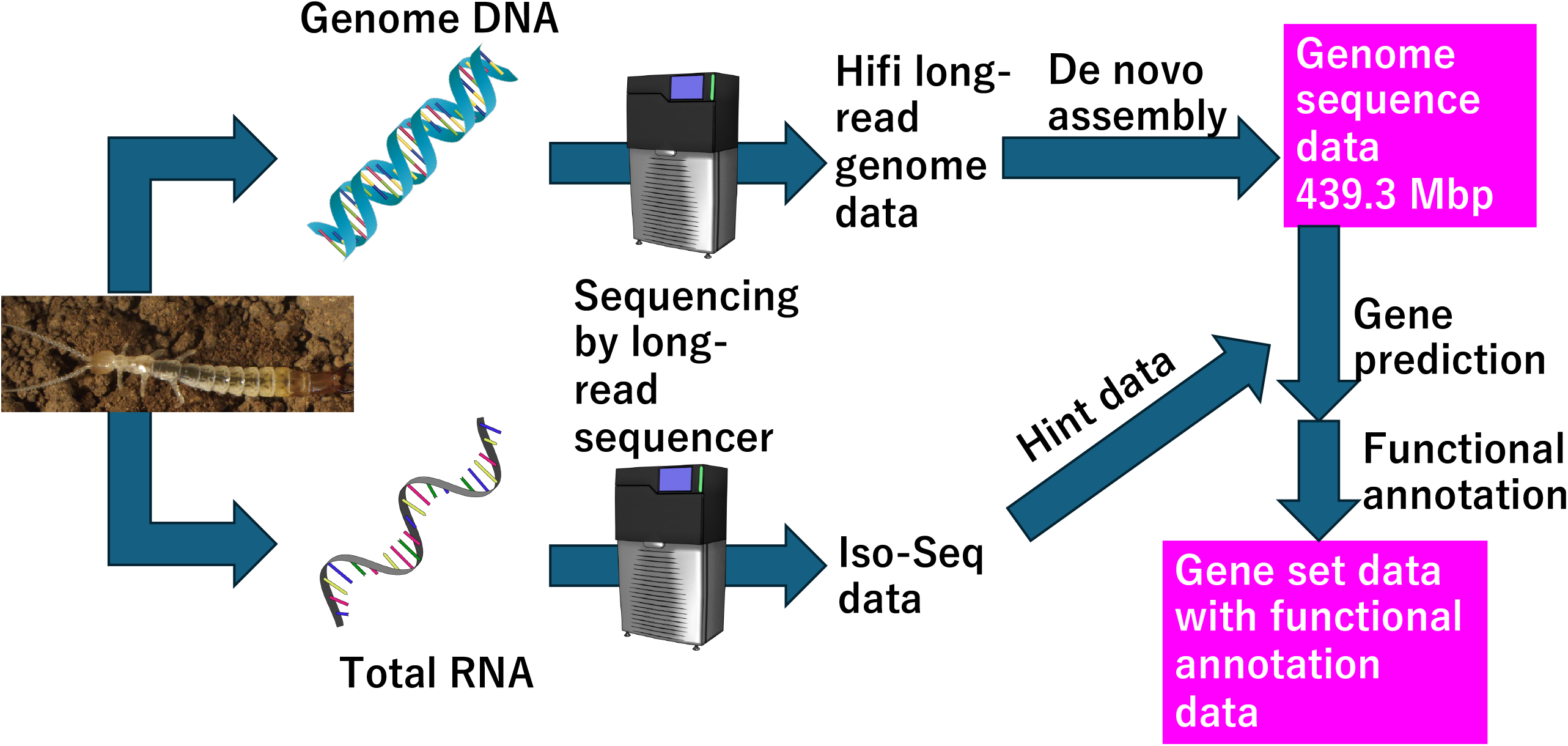
Schematic workflow for the construction of *Occasjapyx japonicus* genome and gene set data. Genomic DNA from a male adult and total RNA from both sexes were extracted and sequenced using the PacBio Revio platform. HiFi long-read genome data and Iso-Seq transcriptomic data were obtained from DNA and RNA, respectively. HiFi reads were used for de novo genome assembly (439.3 Mbp), whereas Iso-Seq data were employed as hints for gene prediction. The resulting gene set was used for functional annotation. Additional data analysis procedures are detailed in Fig. 2.

Whole-genome shotgun sequencing was performed using both PacBio and Illumina sequencing platforms. Genomic DNA was extracted from a male *O. japonicus* specimen using the Genomic-tip Kit (QIAGEN, Hilden, Germany) and sheared with a g-TUBE device (Covaris Inc., MA, USA). DNA fragments shorter than 3 kb were removed using the 35% AMPure PB beads (Pacific Biosciences, CA, USA). A SMRTbell library for high-fidelity (HiFi) sequencing was prepared with the SMRTbell Express Template Prep Kit 3.0 (Pacific Bioscience, CA, USA) according to the manufacturer’s instructions. One SMRT Cell 8M was sequenced on the PacBio Sequel II system using the Sequel II Binding Kit 3.2 and Sequencing Kit 2.0, with 30-hour movie collection time. In addition, a paired-end library was constructed using the Illumina DNA PCR-Free Prep, Tagmentation Kit (Illumina, CA, USA) and an IDT for Illumina DNA/RNA Unique Dual Indexes (Illumina, CA, USA) according to the manufacturer’s instructions. The final library was sequenced on the Illumina NovaSeq 6000 system with the SP Reagent Kit v1.5 (150 bp paired-end mode), generating 297.3 million read pairs per library.

### Data analysis

The data analysis workflow for constructing the *O. japonicus* genome assembly and gene set data is illustrated in Fig. 3. Hifi long-read genome sequence data generated using PacBio Sequel II were assembled with Hifaism (version v0.19.8-r603) to construct the *O. japonicus* genome[14–16]. Assembly metrics were calculated using Assembly-stats (version 1.0.1), and genome quality was assessed using BUSCO (version 5.2.2) against the arthropoda_odb10 database[17]. The genome size prediction was done by the GenomeScope 2.0 website (URL: http://genomescope.org/genomescope2.0/)[18] with default settings using k-mer histgram data as input data which are performed by jellyfish version 2.2.1[19]. The genome completeness was assessed by Merqury version 1.3[20].

**Fig. 3.**
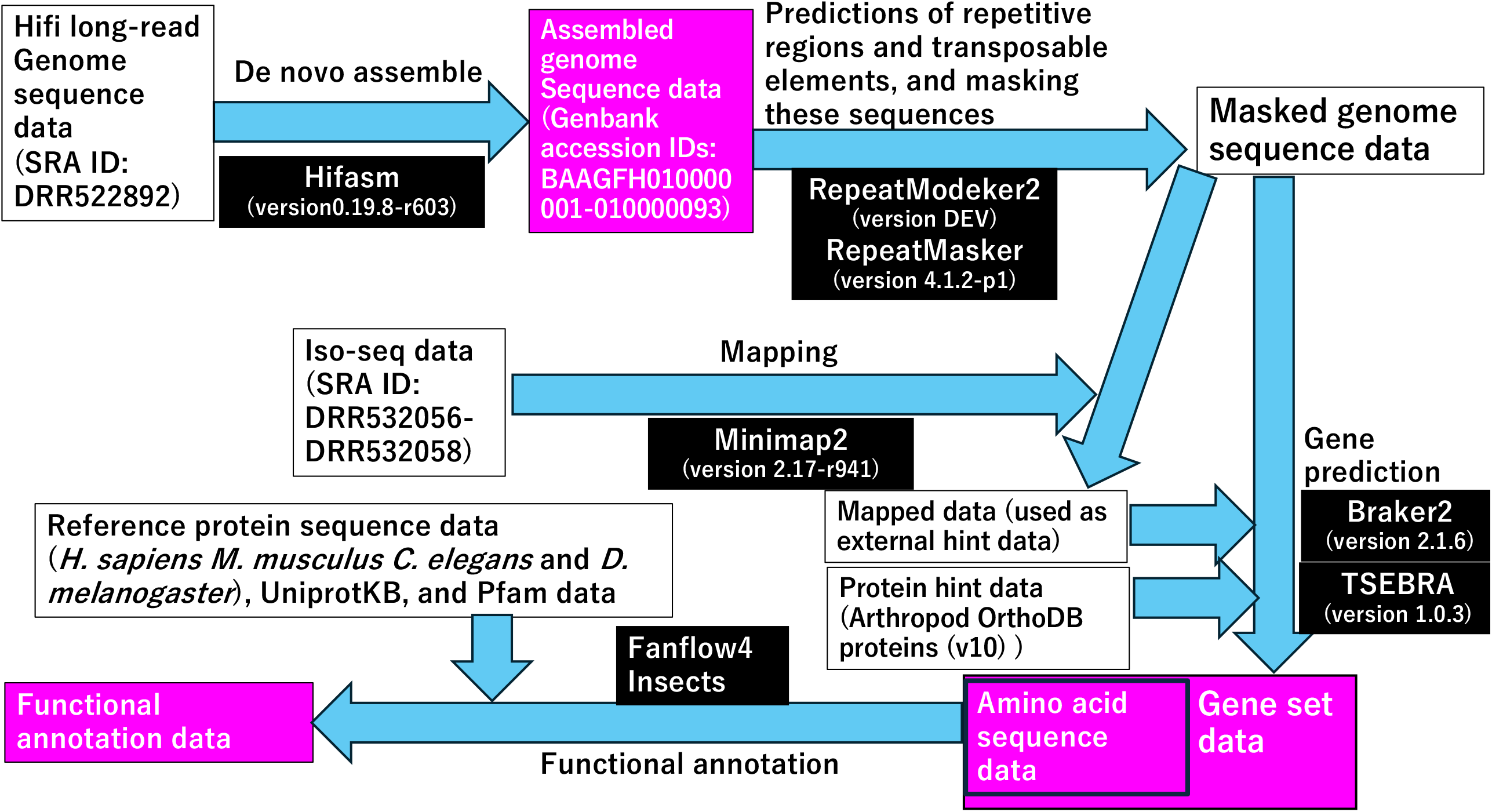
Detailed workflow of the data analysis conducted in this study. White text in purple boxes represents the assembled genome sequence, gene set, and functional annotation data of *Occasjapyx japonicus*. White text in black boxes denotes the software (with version) used for each step of the analysis.

To predict repetitive regions and transposable elements in the *O. japonicusgene* genome, RepeatModeler2 (version DEV)[21] and RepeatMasker (version 4.1.2-p1)[22] were used. The masked genome sequences data in which sequences (with repeats substituted by “N”) were then used as input for gene prediction.

Two types of hint data (Iso-seq and OrthoDB protein datasets) were prepared to generate the *O. japonicusgene* gene set. The first was a BAM file containing Iso-Seq reads mapped to the assembled genome. Mapping was conducted using Minimap2 (version 2.17-r941)^21^, and SAM files were converted to the BAM format using Samtools (version 1.1). The second dataset was the Arthropod OrthoDB protein file (v10), obtained from OrthoDB[24]. Gene prediction was conducted independently using Braker2(version 2.1.6) [25], with the masked genome and either Iso-Seq BAM data or OrthoDB protein data as hints. The two resulting gene prediction datasets were merged using TSEBRA version 1.0.3[26]. The merged dataset constituted the final gene set. Functional annotations was conducted using the predicted amino acid sequences as queries against reference protein datasets from model organisms, including *Homo sapiens* (version GRCh38; Homo_sapiens.GRCh38.pep.all.fa.gz)[27], *Mus musculus* (version GRCm39; Mus_musculus.GRCm39.pep.all.fa.gz)[28], *Caenorhabditis elegans* (version WBcel235; Caenorhabditis_elegans.WBcel235.pep.all.fa)[29], *Drosophila melanogaster* (Drosophila_melanogaster.BDGP6.32.pep.all.fa.gz)[30], Unigene data (Uniprot KB proteint data, uniprot_sprot.fasta.gz), and Pfam data (Pfam-A.hmm), using the Fanflow4Insects[31].

### Data records and code availability

Raw sequence data have been deposited in the DDBJ database (Bioproject No. PRJDB16939, the Sequence Read Archive (SRA) ID: DRR532056-DRR532058 (Full-Length Non-Chimeric), DRR522891, and DRR522892). The assembled genome is available in GenBank (ID: BAAGFH010000001-010000093). Tables listing Contig and GenBank IDs have been uploaded to Figshare[32]. The results of repetitive region and transposable element predictions in *O. japonicusgene*, generated using ReperatModeler2 and RepeatMasker, are available on Figshare[33]. Gene set and functional annotation data have been deposited in Figshare[34]. The code file and associated scripts used for the data analysis in this study (Fig. 2) are available on Figshare[35].

## Results and Discussion

The assembled genome comprises 93 contigs, with a total size of approximately 439 Mbp and an N50 value of 15.5 Mbp (Table 1). The CG content is 41.40%, with average contig lengths of approximately 4.8 Mbp and a maximum contig size of 47.3 Mbp. Repetitive regions and transposable elements constitute 44.14% of the *O. japonicus* genome. A total of 12,628 retrotransposable elements (1.09%) and 61,288 DNA transposons (4.24%) were identified (see detailed results in Figshare)[33]. A total of 19,635 protein-coding genes were predicted in the *O. japonicus* genome (Table 1). As described in the “Technical Validation” section, the genome and the gene set data are of high quality and reliability as reference datasets. These data provide a foundation for exploring the genetic and molecular characteristics of Diplura. Furthermore, these data can be utilized to facilitate comparative studies across species, offering novel evolutionary insights, particularly into hexapod evolution and the emergence of insects.

**Table 1.** Basic status of *Occasjapyx japonicusgene* genome.

|  |  |
| --- | --- |
| Total length | 439,367,470 bp |
| Number of contigs | 93 |
| Average length of the contigs | 4,724,381.40 bp |
| Largest size among the contigs | 47,264,139 bp |
| N50 | 15,511,673 |
| CG contains | 41.40% |
| Number of the predicted genes | 19,635 |

To validate the assembled genome and gene set, we performed the following assessments. BUSCO software assesses genome quality by identifying core genes (“BUSCOs”) that are conserved across a wide range of species[17]. As shown in Table 2, 99% of the BUSCO genes were identified as complete sequences. Approximately 96.9% of BUSCOs were identified as single-copy genes. These BUSCO results indicate that the assembled genome is of high quality in terms of core gene completeness, supporting its use as a reference genome.

**Table 2.** BUSCO Results Using arthropoda_odb10 database (Total 1013 BUSCOs)

| BUSCO Category | Total Numer of BUSCOs | Ratio (Identified BUSCOs/Total BUSCOs) |
| --- | --- | --- |
| Complete BUSCOs | 1003 | 99% |
| Complete and single-copy BUSCOs | 982 | 96.90% |
| Complete and duplicated BUSCOs | 21 | 2.10% |
| Fragmented BUSCOs | 5 | 0.5% |
| Missing BUSCOs | 5 | 0.5% |

The genome assembly was generated using raw long-read sequencing data. The resulting N50 value exceeded 15 Mb, which is significantly higher than that reported for other dipluran genome assemblies (Table 3) and also exceeds that of long-read genome assemblies from other species (e.g., *Mythimna separata*; N50 = 2.7 Mb). These results demonstrate that the assembled genome is of high quality in terms of contiguity.

**Table 3.** Comparative analysis of N50 values, gene counts, and genome sizes across dipluran.

| Species (Accession IDs of GenBank) | *Occasjapyx japonicus*<br>(this study) | *Campodea augens*<br>(GCA_009757345.1) | *Catajapyx aquilonaris*<br>(GCA_000934665.2) | *Campodea plusiochaeta*<br>(GCA_034698565.1) | *Campodea silvestrii*<br>(GCA_034694985.1) |
| --- | --- | --- | --- | --- | --- |
| N50 (contig or scaffold (if exist)) | 15.5 Mbp | 235.2 kbp | 45.9 kbp | 3.6 kbp | 2.5 kbp |
| Genome size | 439.3 Mbp | 1.1 Gbp | 285.6 Mbp | 562.7 Mbp | 776.1 Mbp |
| Number of genes (protein coding) | 19635 | 23992* | 10901*,** | N/A | N/A |

\* Manni et al., (2019) DOI:10.1093/gbe/evz260
\*\* https://i5k.nal.usda.gov/Catajapyx_aquilonaris

The genome size of *O. japonicus* was compared with those of four other diplurans species (*C. augens, C. aquilonaris, C. plusiochaeta*, and *C. silvestrii*) (Table 3). Although the genome of *C. augens* (1.1 Gbp) is larger than that of *O. japonicus*, the genome sizes of the other three species are comparable to, or not significantly different from, that of *O. japonicus*.

Genome size prediction of *O. japonicus* using short-read genome data is performed by Genome Scope. The predicted size of approximately 339 Mbp, which is about 100 Mbp different from the size of the actual assembled genome data. The model precision values and error rate of the GenomeScope were about nearly zero and 0.5%, respectively, suggesting that there is no critical error in the GenomeScope analyses. Also, output graphs from the GenomeScope showed that the precition is reliable (Fig. 4). As shown above, 44.14% of the *O. japonicus* genome consists of repetitive and TE regions, which are not precisely assembled using the short-read genome data. The GenomeScope size prediction was performed using short-read data. Considering them, these differences in the genome size between the prediction and the actual assembled genome data might be due to these repetitive and TE regions. Furthermore, the genome quality evaluation was performed by Merqury. The QV value and completeness ratio of the assembled genome are 55.9144 and 97.5813%, respectively, suggesting that the genome data possess sufficient quality for a reference genome. The assembled gene set for *O. japonicus* included 19,635 protein-coding genes. *Campodea augens* and *Catajapyx aquilonaris* contain 23,992 and 10,901 protein-coding genes, respectively. Thus, the number of protein-coding genes in *O. japonicus* falls within the range observed in these two species (Table 3).

**Fig. 4.**
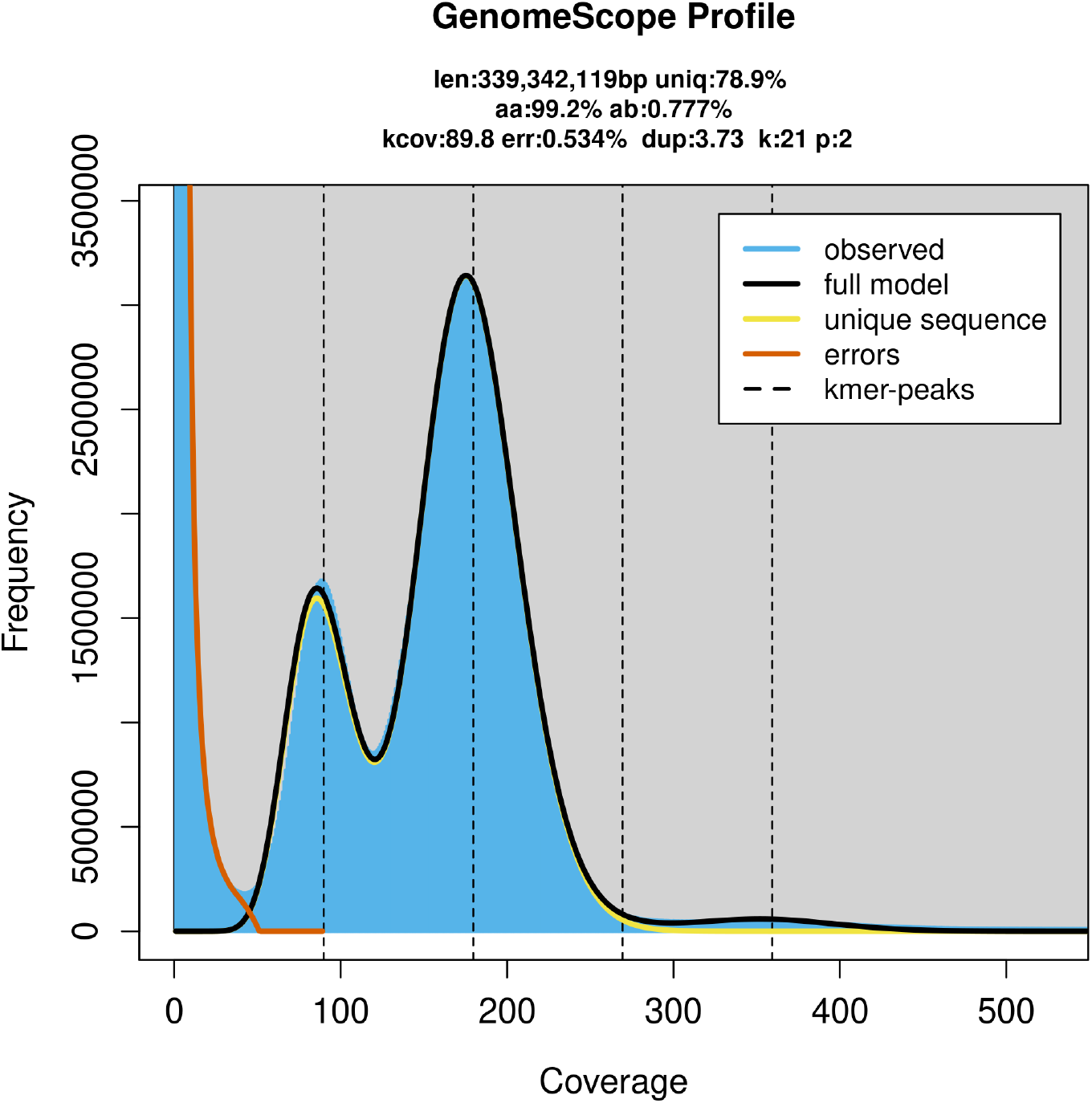
Genome Scope output graph.

In conclusion, the results described above indicate that the genome assembly and gene set for *O. japonicus* are of sufficient quality and reliability to serve as reference genomic resources.

## Acknowledgements

This work was supported by JSPS KAKENHI (Grant Numbers 23K26918 and JP22H04925 [PAGS]) awarded to K.Y., K.H., and T.A.

We express our gratitude to Drs. Akiya Jouraku and Kohei Toga for advice on the data analyses.

Parts of Figs. 2 and 3 were drawn using illustrations from TogoTV (© 2016 DBCLS TogoTV, CC-BY-4.0 https://creativecommons.org/licenses/by/4.0/deed.ja ).

## Author contributions

K.Y. and T.A. conceived the study. T.A. prepared DNA and RNA samples from *O. japonicus*. K.Y. performed bioinformatics data analysis and data registration. A.T. obtained the sequencing data. K.H. identified the species used in the analysis. K.Y. and T.A. wrote the original draft of the manuscript, and reviewed and edited the draft of the manuscript. All authors have read and agreed to the published version of the manuscript.

## References

1. Schwentner, M., Combosch, D. J., Pakes Nelson, J. & Giribet, G. A Phylogenomic Solution to the Origin of Insects by Resolving Crustacean-Hexapod Relationships. Current Biology 27, 1818-1824.e5 (2017).

2. Giribet, G. & Edgecombe, G. D. The Phylogeny and Evolutionary History of Arthropods. Current Biology 29, R592–R602 (2019).

3. Beutel, R. G., M. I. Yavorskaya, Y. Mashimo, M. Fukui and K. Meusemann. The Phylogeny of Hexapoda (Arthropoda) and the Evolution of Megadiversity. In Proc. Arthropod. Embryol. Soc. Jpn 1–15.

4. Du, S. et al. Revisiting the four Hexapoda classes: Protura as the sister group to all other hexapods. Proceedings of the National Academy of Sciences 121, e2408775121 (2024).

5. Misof, B. et al. Phylogenomics resolves the timing and pattern of insect evolution. Science 346, 763–767 (2014).

6. i5K Consortium. The i5K Initiative: advancing arthropod genomics for knowledge, human health, agriculture, and the environment. J Hered 104, 595–600 (2013).

7. Manni, M. et al. The Genome of the Blind Soil-Dwelling and Ancestrally Wingless Dipluran Campodea augens: A Key Reference Hexapod for Studying the Emergence of Insect Innovations. Genome Biology and Evolution 12, 3534–3549 (2020).

8. NCBI genome Campodea plusiochaeta https://www.ncbi.nlm.nih.gov/datasets/genome/GCA_034698565.1/. (2023).

9. NCBI genome Campodea silvestrii https://www.ncbi.nlm.nih.gov/datasets/genome/GCA_034694985.1/. (2023).

10. Enderlein, G. Über die Segmental-Apotome der Insekten und Kenntnis der Morphologie der Japygiden. Zoologischen Anzeiger 31, 629–635 (1907).

11. Kuwayama, S. Some Japanese Species of Japyx. Insecta Matsumurana 2, 151–155 (1928).

12. Silvestri, F. Japygidae (Thysanura) dell’Estremo Oriente. Bollettino del Laboratorio di zoologia generale e agraria della R. Scuola superiore d’agricoltura in Portici 22, 49–8 (1928).

13. Pagés, J. Sclérites et appendices de l’abdomen des Diploures (Insecta, Apterygota). Archives des Sciences Genève 42, 509–551 (1989).

14. Cheng, H., Concepcion, G. T., Feng, X., Zhang, H. & Li, H. Haplotype-resolved de novo assembly using phased assembly graphs with hifiasm. Nat Methods 18, 170–175 (2021).

15. Cheng, H. et al. Haplotype-resolved assembly of diploid genomes without parental data. Nat Biotechnol 40, 1332–1335 (2022).

16. Cheng, H., Asri, M., Lucas, J., Koren, S. & Li, H. Scalable telomere-to-telomere assembly for diploid and polyploid genomes with double graph. Nat Methods 21, 967–970 (2024).

17. Manni, M., Berkeley, M. R., Seppey, M., Simão, F. A. & Zdobnov, E. M. BUSCO Update: Novel and Streamlined Workflows along with Broader and Deeper Phylogenetic Coverage for Scoring of Eukaryotic, Prokaryotic, and Viral Genomes. Molecular Biology and Evolution 38, 4647–4654 (2021).

18. Ranallo-Benavidez, T. R., Jaron, K. S. & Schatz, M. C. GenomeScope 2.0 and Smudgeplot for reference-free profiling of polyploid genomes. Nat Commun 11, 1432 (2020).

19. Marçais, G. & Kingsford, C. A fast, lock-free approach for efficient parallel counting of occurrences of k-mers. Bioinformatics 27, 764–770 (2011).

20. Rhie, A., Walenz, B. P., Koren, S. & Phillippy, A. M. Merqury: reference-free quality, completeness, and phasing assessment for genome assemblies. Genome Biol 21, 245 (2020).

21. Flynn, J. M. et al. RepeatModeler2 for automated genomic discovery of transposable element families. Proc Natl Acad Sci U S A 117, 9451–9457 (2020).

22. Smit, A., Hubley, R. & Green, P. RepeatMasker Open-4.0. http://www.repeatmasker.org (2013).

23. Li, H. Minimap2: pairwise alignment for nucleotide sequences. Bioinformatics 34, 3094–3100 (2018).

24. Kriventseva, E. V. et al. OrthoDB v10: sampling the diversity of animal, plant, fungal, protist, bacterial and viral genomes for evolutionary and functional annotations of orthologs. Nucleic Acids Research 47, D807–D811 (2019).

25. Brůna, T., Hoff, K. J., Lomsadze, A., Stanke, M. & Borodovsky, M. BRAKER2: automatic eukaryotic genome annotation with GeneMark-EP+ and AUGUSTUS supported by a protein database. NAR Genomics and Bioinformatics 3, lqaa108 (2021).

26. Gabriel, L., Hoff, K. J., Brůna, T., Borodovsky, M. & Stanke, M. TSEBRA: transcript selector for BRAKER. BMC Bioinformatics 22, 566 (2021).

27. NCBI genome Hono sapiense https://www.ncbi.nlm.nih.gov/datasets/genome/GCF_000001405.40/. (2022).

28. NCBI genome Mus musculus https://www.ncbi.nlm.nih.gov/datasets/genome/GCF_000001635.27/. (2020).

29. NCBI genome Caenorhabditis elegans https://www.ncbi.nlm.nih.gov/datasets/genome/GCF_000002985.6/. (2013).

30. Ensembl genome Drosophila melanogaster https://feb2023.archive.ensembl.org/Drosophila_melanogaster/Info/Index. (2020).

31. Bono, H., Sakamoto, T., Kasukawa, T. & Tabunoki, H. Systematic Functional Annotation Workflow for Insects. Insects 13, 586 (2022).

32. Yokoi, K., Toyada, A., Hashimoto, K. & Asano, T. List of Contig IDs and Genbank accession IDs. Figshare 10.6084/m9.figshare.28262774 (2025).

33. Kakeru, Y., Atsushi, T., Kosei, H. & Tsunaki, A. The result data of prediction of repetitive regions and transposable elements in O. aponicusgene. Figshare 10.6084/m9.figshare.28262804 (2025).

34. Yokoi, K., Toyoda, A., Hashimoto, K. & Asano, T. Occasjapyx japonicus gene set data and functional annotation data of the gene set. Figshare 10.6084/m9.figshare.28263434 (2025).

35. Yokoi, K., Toyoda, A., Hashimoto, K. & Asano, T. Script files for data analysis. Figshare 10.6084/m9.figshare.28263503 (2025).

